# Post-zygotic reproductive isolation is not correlated with chromosome number in plants

**DOI:** 10.1101/2024.01.05.573914

**Authors:** Geoffrey S Finch, Yang Yang, Michael S Barker

## Abstract

The evolution of chromosome numbers is an important but not fully understood aspect of eukaryotic evolution. Although we understand the types of karyotypic changes that can lead to chromosome gain and loss, we still do not understand why chromosome numbers in many plants and animals have an average of *n* = 9. A recent hypothesis proposed that chromosome number reduction following whole genome duplication (WGD) in angiosperms is driven by an interaction between chromosome number and the strength of reproductive isolation among populations. Chromosome number is expected to determine the maximum number of independently assorting Bateson-Dobzhansky-Muller incompatibilities (BDMIs). Selection against restricted gene flow among populations would result in selection for reduced chromosome number. Here we test for an interaction between chromosome number and post-zygotic reproductive isolation across a broad sample of land plants. We additionally tested for the indirect effects of WGD in generating post-zygotic reproductive isolation. Such an effect is expected if reproductive isolation is driven largely by reciprocal gene loss of WGD-derived paralogs. We found no correlation between post-zygotic reproductive isolation and chromosome number, WGD age, or degree of fractionation, suggesting that the accumulation of genic incompatibilities is likely not a major driver of post-WGD chromosome number reduction.

## INTRODUCTION

The Bateson-Dobzhansky-Mueller (BDM) model for the evolution of reproductive incompatibilities is an important model of speciation [1–4]. In this model, post-zygotic reproductive isolation arises when new alleles evolve at epistatically interacting loci in different populations. In some cases these new alleles will interact poorly and reduce the fitness of hybrids. The accumulation of these BDM Incompatibilities (BDMIs) is thought to be an important source of post-zygotic reproductive isolation [1–4]. Similarly, reciprocal gene loss (RGL) can occur following gene and whole-genome duplications (WGD). In the RGL model, different populations lose alternate loci through fractionation. When individuals from these populations hybridize, a fraction of their offspring are completely missing a copy of that gene. This is expected to reduce fitness in a manner similar to BDMIs [5–7]. A common element of these models of post-zygotic reproductive isolation is that the fitness effect is strongest when pairs of interacting genes are unlinked. Thus, we might expect that species with more opportunity for unlinked BDMI and RGL sets will experience greater reproductive isolation. One simple way to estimate the potential for unlinked loci is the chromosome number of a species [6]. All else being equal, species with more chromosomes will have more opportunity for unlinked BDMI sets.

An interesting implication of this potential relationship is that selection acting on levels of reproductive isolation between demes or species could influence patterns of chromosome number variation [6]. In angiosperms, haploid chromosome number varies from *n* = 2 to *n* ∼320 [8–11], yet the majority of plants have chromosome numbers around *n* = 9 [11,12]. One well-established pattern of chromosome number evolution in angiosperms is the cycle of chromosome number doubling resulting from WGD and the subsequent reduction in chromosome number during the process of diploidization [13]. This chromosome number reduction is accompanied by the rapid loss of paralogs (i.e. fractionation) which has the potential to create large numbers of RGL sets across the genome [5,6,13,14]. Because of this, neopolyploids with low chromosome numbers may be more successful than species with high chromosome numbers in maintaining gene flow across isolated populations, increasing their odds of survival. This could produce the observed pattern of chromosome number loss seen in angiosperms [6].

We extend this thinking to BDMIs in general, predicting that, as such loci accumulate, closely related species with higher chromosome numbers should experience more pronounced reductions in fitness than those with lower chromosome numbers. In this way, chromosome number is expected to interact with BDMIs to impact fitness regardless of WGD history. To test this hypothesis, we could look for a positive relationship between BDMI sets and chromosome number. However, developing a complete survey of BDMIs for multiple species with varying chromosome numbers is not yet practical. An alternative inference can be made using crossability data in place of BDMIs. If BDMIs are correlated with chromosome number, then by extension we may observe that post-zygotic reproductive isolation in general is correlated with chromosome number. Here, we leverage crossability data from 101 congeneric species pairs or groups of plants to assess the relationship between chromosome number and post-zygotic reproductive isolation. We tested for broad-scale effects of chromosome number on the crossability of closely related species with varied histories of WGD, using a measure of postzygotic reproductive isolation, the crossability index, that is expected to be impacted by the number of independently assorting BDMIs. If RGL is the primary source of reproductive isolation, we may only see an interaction between chromosome number and crossability in taxa with a more recent shared WGD or with a large number of retained paralogs. To account for this possibility, we examined the importance of the age of the most recent WGD shared by a pair of species by testing for correlations between crossability and (1) paralog median Ks, a rough measure of the age of the WGD, and (2) paleolog fraction, a measure of the degree of fractionation that has occurred following WGD [13,15]. These metrics provide a rough idea of how many duplicated genes may still act as sources of RGL in the species examined here. By testing these predictions, we provide new insight into the relationship among chromosome number, BDMI accumulation, and WGD history in driving reproductive isolation in plants.

## MATERIALS AND METHODS

### Taxon sampling and data acquisition

Data on crossability and chromosome numbers were taken from previous studies [11,16]. Crossability indices were obtained from Rieseberg et al. [16]. This is a measure of post-zygotic reproductive isolation defined as the ratio of interspecific crossability to intraspecific crossability for a pair or group of congeneric species. These estimates include only measures of hybrid sterility, such as pollen viability, seed set, or spore viability of the F1. As such, these data are well-suited to test for an interaction between post-zygotic reproductive isolation and chromosome number. The crossability index ranges from zero (complete isolation) to one (total crossability), and values lower than ∼0.8 were considered to have significantly lower interspecific than intraspecific crossability (Rieseberg et al. 2006). Haploid chromosome numbers were obtained from the Chromosome Counts Database [11] for the species used in the crossability studies. We calculated the mean and variance in chromosome number for each genus using the chromosome numbers of the species used to determine crossability indices. Variance was calculated to determine if crossability is determined primarily by chromosome structural differences between taxa, such as chromosome number changes. If so, we may see a reduction in crossability in those species pairs/groups with greater variability in chromosome number.

To test the relationship among WGD age and other variables, we identified the most recent WGD shared by the species for which crossability data were available. The age of a WGD can be approximated by measuring the synonymous divergence (Ks) of paralogs resulting from the duplication, with more divergence among relatively older WGDs. We identified the most recent putative WGD and the median Ks values from previous analyses (1KP, [15,17]). When possible, we used median Ks values for species (n = 14) in the crossability data. If unavailable, the median Ks estimate from a congeneric species (n = 23) or member of a closely related genus (n = 63) was used (Supplemental Table S2).

We also evaluated whether the degree of fractionation is correlated with chromosome number and post-zygotic reproductive isolation among congeneric species. Fractionation is the loss of paralogs following polyploidization as the genome returns to a diploid state (i.e. diploidization). Paralogs resulting from an ancient WGD are called paleologs. We used estimates of the proportion of the genome that has been retained in duplicate following a WGD in each of these lineages—the paleolog fraction—from a previous study [13]. If paleolog fraction estimates for multiple members of a genus were available, the mean value for the genus was used (Supplemental Table S2).

Sample sizes for each analysis depended on the availability of estimates for median Ks, paleolog fraction, and crossability index, as well as chromosome counts and *rbcL* sequences (see below). In total, crossability indices from 98 genera were used in our analyses: In the full data set, 97 samples (i.e. pairs/groups of species used to estimate crossability index) were available for regressions on chromosome number, 99 samples for median Ks, and 31 samples for paleolog fraction. To account for potentially different patterns of diploidization in angiosperms and other plant clades [18], we further divided the dataset into ‘angiosperms’ and ‘non-angiosperms.’

### Regression analyses

To account for phylogenetic relationships in our analyses, we used phylogenetic generalized least squares (PGLS) to test for correlations between crossability index and mean haploid chromosome number, variance in chromosome number, median Ks, and paleolog fraction. A species tree was constructed using *rbcL* sequences from genBank and aligned in PASTA (1.7.8) [19]. The list of genBank accession IDs and associated species names for the *rbcL* sequences used is available in Supplemental Table S1. An ultrametric tree was constructed using RAxML (8.2.11) [20] under a GTR + gamma model of sequence evolution and the *chronos* function in the R package ape (5.6.2) [21]. As the regression analyses only use information from relative node ages, the root age was arbitrarily set to 1. PGLS was conducted using the *corPagel* variance structure with the R function *gls*, part of the ape package. The *gls* function with the *corPagel* variance structure simultaneously estimates regression coefficients as well as the phylogenetic signal in the residual errors, represented by lambda (λ) [21,22].

## RESULTS

Using data from 99 genera of land plants, we found no relationship between crossability and chromosome number as would be expected if higher chromosome numbers led to more independent BDMI sets. The PGLS analysis of crossability and chromosome number across the 99 genera (500 species and subspecies) was not significant with no trend apparent (β = 0.00013, p = 0.96, λ = 0.41, n = 97; Figure 1a). Because angiosperms have relatively high rates of diploidization and chromosome number evolution compared to many non-angiosperm clades [13] we also tested for a relationship just in angiosperms and non-angiosperms. Similar to our overall analyses, we did not find a significant correlation between crossability and chromosome number in either group (angiosperms: β = 0.0008, p = 0.88, λ = 0.25, n = 85; Figure 1b; non-angiosperms: β = 0.0009, p = 0.62, λ = -0.16, n = 12; Figure 1c). These results suggest that postzygotic reproductive isolation in these clades is not determined by chromosome number (Table 1). If crossability is determined primarily by chromosome structural differences between taxa, such as chromosome number changes, we may see a reduction in crossability in those species pairs/groups with greater variability in chromosome number. To test this possibility, we also tested for a correlation between crossability index and variance in chromosome number. No relationship was detected in any subset of the data (all species: β = 0.00008, p = 0.81, λ = 0.45, n = 85; angiosperms: β = -0.0003, p = 0.76, λ = 0.29, n = 85; non-angiosperms: β = 0.00001, p = 0.95, λ = -0.46, n = 12).

**Table 1.** Summary of results of PGLS regressions. From left to right, the slope of the regression line (β), p-value, estimate of phylogenetic signal (λ), and sample size are shown for each dataset. All results are non-significant. Graphs of corresponding data and regression lines are shown in Figure 2.

|  | All species |  |  |  | Angiosperms |  |  |  | Non-angiosperms |  |  |  |
| --- | --- | --- | --- | --- | --- | --- | --- | --- | --- | --- | --- | --- |
| | $\beta$ | p | $\lambda$ | n | $\beta$ | p | $\lambda$ | n | $\beta$ | p | $\lambda$ | n |
| Chromosome Number | 0.00013 | 0.96 | 0.410 | 97 | 0.00081 | 0.88 | 0.25 | 85 | 0.00031 | 0.83 | -0.41 | 12 |
| Median Ks | 0.014 | 0.85 | 0.350 | 99 | 0.029 | 0.72 | 0.29 | 85 | 0.04 | 0.80 | -0.18 | 14 |
| Paleolog Fraction | -0.13 | 0.87 | -0.057 | 35 | 0.061 | 0.95 | 0.40 | 24 | 0.55 | 0.69 | -0.36 | 11 |

**Figure 1.**
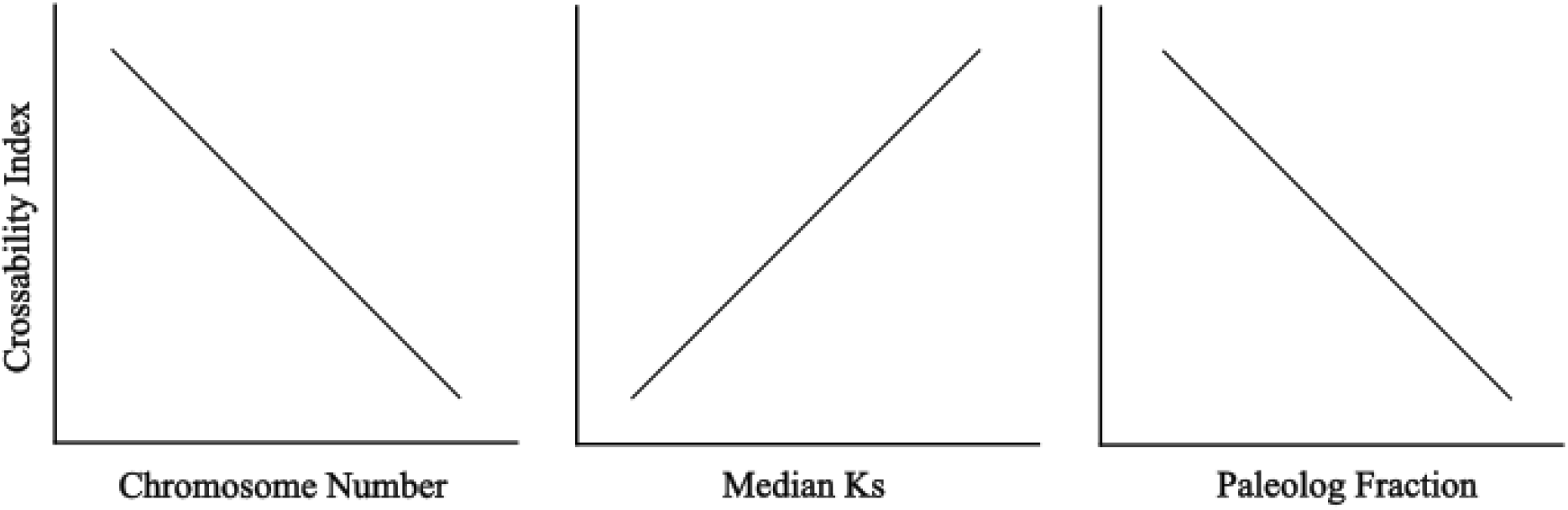
Summary of expected relationships between crossability index and (left) chromosome number if higher chromosome number results in higher numbers of independently assorting BDMIs, (center) paleolog median Ks if more recent shared WGDs result in higher numbers of independently assorting RGL sets, and (right) paleolog fraction if higher proportions of paleologs result in higher numbers of independently assorting RGL sets. These are simplified representations of expectations, and trends are not necessarily linear. Increasing chromosome number is predicted to decrease crossability as a result of the increase in the number of independently assorting BDMI sets. Median Ks is a measure of the age of a shared WGD. Based on the hypothesis that RGL results in reproductive isolation in diploidizing lineages, we expect crossability to decrease in those taxa with a relatively recent shared WGD. Similarly, taxa with high numbers of paleologs are expected to experience reduced crossability.

Although we find no evidence that chromosome number variation in general is associated with crossability, Bowers and Paterson (6) specifically proposed that an increase in RGL sets following WGD may drive reductions in chromosome number. If this is the case we expect to see an effect of WGD history on levels of reproductive isolation. Specifically, a recent shared WGD should result in reduced crossability among descendant species (Fig. 1). To test this hypothesis more directly, we examined the relationships between shared WGD age and degree of fractionation on crossability. First, we tested if there was a relationship between the divergence of WGD paralogs and crossability. If reciprocal gene loss (RGL) following WGD is an important contributor to post-zygotic reproductive isolation in plants, we expect a positive correlation between paralog divergence and crossability, because genera with more recent WGDs may have more opportunity for RGL to drive reproductive isolation (Fig. 1). We collected median Ks estimates for 35 WGDs in the ancestry of these lineages, with median Ks ranging from 0.1733 to 2.6110. Paralog divergence and crossability index were not significantly correlated across all taxa (β = 0.014, p = 0.85, λ = 0.35, n = 99; Fig. 2d), or when we separated angiosperms (β = 0.029, p = 0.72, λ = 0.29, n = 85; Fig. 2e) and non-angiosperms (β = 0.00006, λ = -0.15, p = 0.9997, n = 12; Fig. 2f). Thus, paralog divergence, as represented by median Ks, is not a strong predictor of the level of reproductive isolation in the taxa analyzed here (Table 1). That is, reproductive isolation does not appear to accumulate more rapidly in taxa with a more recent shared WGD, and this pattern does not differ between angiosperms and non-angiosperms.

**Figure 2.**
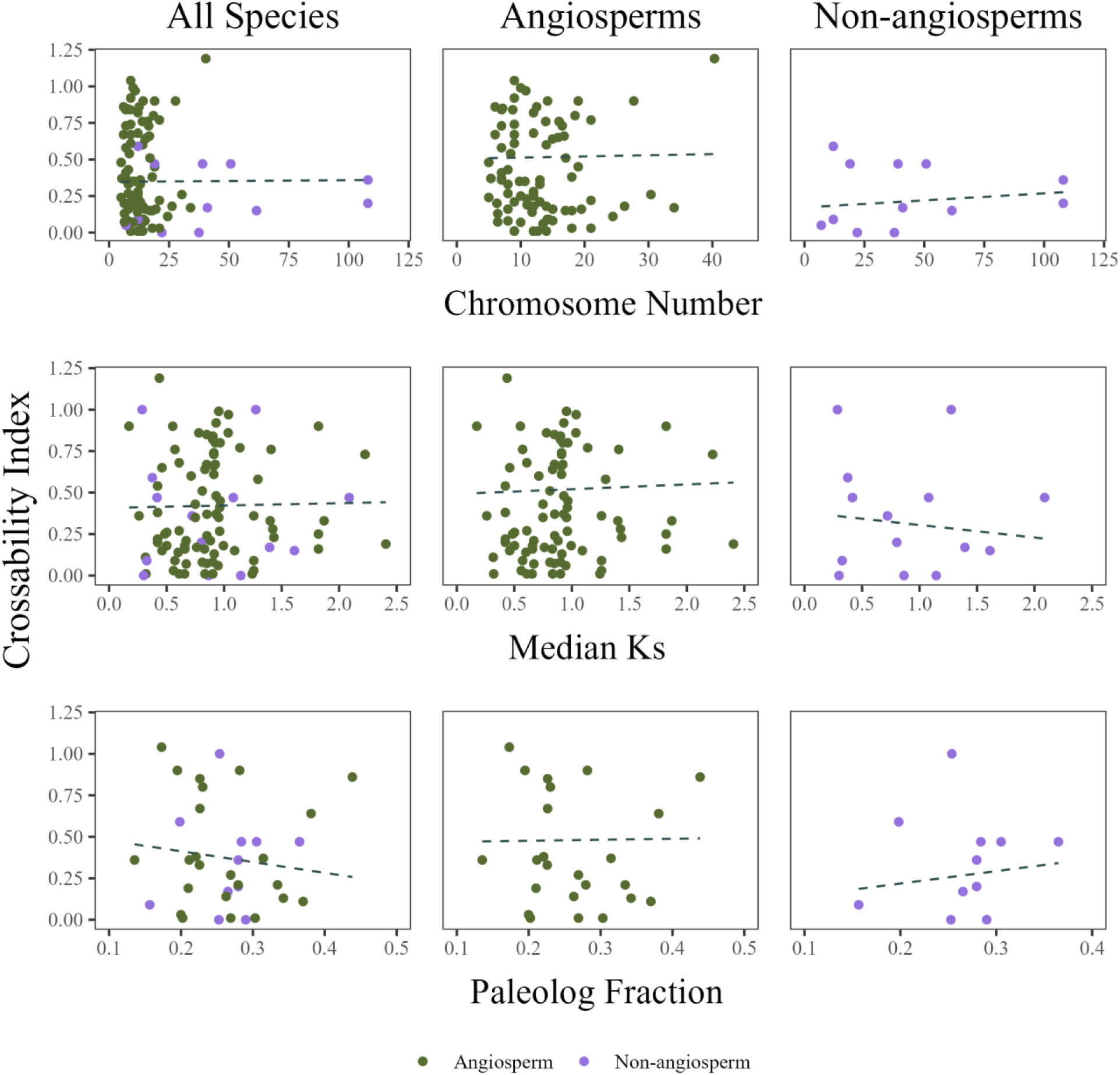
Results of PGLS regressions of crossability index (CI) on haploid chromosome number (top), paleolog median Ks (middle), and paleolog fraction (bottom) for all species (left), angiosperms (middle, green), and non-angiosperms (right, purple). No significant relationship is detected for any variable in any dataset.

Our second variable associated with WGD history, paleolog fraction, is an estimate of the proportion of the genome that has been retained in duplicate following a WGD. It is the number of paleologs divided by the number of unigenes in the transcriptome and is one measure of how much fractionation has occurred in the genome [13]. Species with relatively high paleolog fractions have the potential for more RGL to occur among congeneric species and putatively drive reproductive isolation. Because gene loss is the driver of reproductive isolation under the reciprocal gene loss model for post-polyploid diversification [5,6], paleolog fraction and crossability index are expected to be negatively correlated (Fig. 1). Similar to our previous analyses, we did not find evidence for the predicted relationship between crossability and paleolog fraction. Paleolog fraction ranged from 0.1354 to 0.4384 across 80 taxa. There was no relationship between crossability index and paleolog fraction across all taxa (β = 0.20, p = 0.80, λ = -0.05, n = 31; Figure 2g), or when just comparing among angiosperms (β = 0.26, p = 0.79, λ = 0.166, n = 21; Figure 2h) or non-angiosperms (β = -0.079, p = 0.98, λ = -0.20, n = 8; Figure 2i). As with median Ks, these results indicate that reproductive isolation does not accrue more quickly in lineages with a higher proportion of paleologs (Table 1).

## DISCUSSION

We find no evidence that higher numbers of chromosomes are associated with increased post-zygotic reproductive isolation in a diverse collection of 99 plant genera. We detected no relationship between haploid chromosome number and crossability index in any subset of our data. This suggests that, across broad phylogenetic scales, chromosome number is either not a good predictor of the number of independently assorting BDMI sets in a given genome, that BDMIs are not an important determinant of hybrid sterility in plants, or both. The latter could result from a paucity of such loci in general (i.e. genes rarely cause hybrid sterility in plants); alternatively, such loci may not interact with chromosome number in the predicted manner. In either case, chromosome number variation is likely unaffected by selection to reduce the number of independently assorting BDMI sets.

In addition, the crossability of closely related species does not appear to be affected by degree of fractionation or the age of the most recent WGD common to the species in question. We detected no significant relationships between crossability and paleolog fraction or paralog median Ks of WGDs in the ancestry of each genus. WGD-derived RGL sets have been proposed to explain the rapid cytological diploidization and chromosome number reduction observed in angiosperms (Bowers and Paterson, 2021). If this is the case we expect to see an increase in reproductive isolation in lineages with a more recent shared WGD (Fig. 1). Our results are not consistent with this prediction, suggesting that WGD history is not a broadly important determinant of reproductive isolation in plants. Moreover, our observed patterns did not differ between angiosperms and non-angiosperms, two groups that exhibit different patterns of chromosome number evolution following WGD [13]. For example, homosporous ferns appear to experience WGD without the same degree of chromosome loss following WGD as seen in angiosperms, resulting in much higher chromosome numbers overall [11,12,15,23,24]. If WGD-derived RGL sets drive chromosome number reductions in angiosperms only, we expect to see an effect of WGD age on crossability in angiosperms but not other lineages. However, it is not clear why this mechanism would not also drive chromosome number reduction in ferns and gymnosperms. Indeed, our results suggest that these contrasting patterns of diploidization are not somehow driven by differing patterns of RGL or differing responses to the accumulation of RGL sets.

There are many possible explanations for our results. One is that chromosome number is not a good proxy for global recombination rates across the broad phylogenetic scales analyzed here [25,26]. The effect of having fewer chromosomes on the number of independently assorting BDMIs may be very small if meiotic crossovers between BDMI loci are common. In this case, two loci can be freely recombining even if on the same chromosome. Additionally, empirical findings that multiple BDMIs are necessary in some cases to result in negative fitness effects in the hybrid [4] suggest that it may not be straightforward to predict the impacts of changes in chromosome number. In cases of complex epistatic interactions resulting in hybrid sterility, higher chromosome numbers may actually increase gene flow between species by reducing the odds of introgressing both (or all) of the loci involved in the BDMI phenotype [4,27]. Nonetheless, when testing for broad patterns across the phylogenetic scales used here, we can imagine that each introgressed codon with a non-synonymous change has a certain probability of resulting in a negative fitness consequence in the hybrid background, as in [4,28]. As such, the maximum number of independently assorting factors contributing to reproductive isolation is largely set by chromosome number, as described by [6].

If this effect is strictly limited to RGL sets formed during the earliest stages of diploidization, it is possible that our data did not provide the resolution necessary to detect this phenomenon. Although reciprocal gene loss of duplicates derived from small-scale duplications, as well as classic BMDIs, can both result in hybrid sterility in plants [7,29–32], RGL is thought to be especially relevant to neopolyploid lineages [5,6,14]. This is because of the sheer number of gene duplicates that create the opportunity for RGL, and because the number of paralogs declines rapidly following WGD [5,6,13,14,33,34]. The same polyploid lineages can also form independently in multiple locations, potentially giving rise to small, isolated subpopulations that might benefit from reciprocal gene flow [6,35,36]. While there is some evidence that RGL has been associated with speciation following WGD [14,34,35,37], RGL does not appear to be an important driver of speciation in plant clades examined to date [38,39]. We do not at present have the data to directly test whether chromosome number and number of RGL sets interact to determine crossability in only neopolyploid lineages. However, we analyzed a diverse set of taxa with varying histories of WGD, some with substantial numbers of retained paleologs (up to a paleolog fraction of 0.44). In spite of this potential ‘fuel,’ we do not find evidence for a significant interaction between WGD age or degree of fractionation and crossability. These results do not support an outsized role for WGD-derived RGL in determining crossability. Indeed, theory indicates that the conditions under which RGL sets will form are quite limited [33]).

The primary drivers of post-zygotic reproductive isolation among plant species may also limit this mechanism. In contrast to animals, genome structural divergence is thought to be a more common driver of reproductive isolation than genic factors [27]. This has long been supported by the observation that genome doubling in plants commonly restores fertility in hybrids [1,27,40], a result that is unexpected if BDMIs are the primary cause of sterility. Analogous tests for interactions between chromosome number and crossability in animals would provide additional valuable insight into the relative importance of genic sources of hybrid sterility in establishing species boundaries and potentially influencing chromosome number evolution.

## CONCLUSIONS

We find no effect of chromosome number, paleolog fraction, or median Ks on crossability, suggesting that the number of independently assorting BDMI sets in a given plant lineage (regardless of WGD history) is not a major determinant of post-zygotic reproductive isolation. This is consistent with the hypothesis that structural variation is in general a more important driver of RI in plants than genic incompatibilities.

## Supporting information

ultrametric rbcL tree

R code for pgls

## DATA ACCESSIBILITY

All data and code are available in the electronic supplementary materials S1-S4.

## AUTHOR CONTRIBUTIONS

G.S.F and M.S.B. designed research; G.S.F. and Y.Y. performed research and analysed data, and G.S.F. and M.S.B. wrote the paper.

## ACKNOWLEDGEMENTS

We thank Michael McKibben, Justin Conover, Sylvia Kinosian, Jay Goldberg, Qiuyu Jiang, Seongyeon Kang, and Loren Rieseberg for comments and discussion of this manuscript.

